# Effect of cannabinol, tetrahydrocannabivarin and cannabidiol on voluntary alcohol consumption

**DOI:** 10.1101/2025.09.12.675769

**Authors:** Ieva Poceviciute, Martynas Arbaciauskas, Rokas Buisas, Osvaldas Ruksenas, Valentina Vengeliene

## Abstract

Previous studies demonstrated that the endocannabinoid system plays a significant role in the development of the alcohol use disorder (AUD), and CB1 receptor antagonists/inverse agonists showed promise as a novel AUD pharmacotherapy. However, these compounds failed in clinical trials due to the severe psychiatric side effects. It has been suggested that non-psychoactive phytocannabinoids may have a better safety profile and could be used as an alternative approach to treat AUD. Hence, the aim of this study was to test the potential of three phytocannabinoids in reducing alcohol consumption: CB1 receptor partial agonist cannabinol (CBN), neutral antagonist tetrahydrocannabivarin (THCV) and negative allosteric modulator cannabidiol (CBD). For this purpose, male Wistar rats were subjected to a long-term voluntary alcohol drinking procedure that lasted for at least 6 months. Thereafter, rats were given 3 daily administrations of CBN, THCV or CBD. A side-effect profile of all three compounds was measured by recording water consumption, body weight changes and home-cage locomotor activity. Ultrasonic vocalisations were recorded before and after a single administration of CBN, THCV or CBD in alcohol-naïve group-housed rats to monitor if these compounds induced discomfort, distress or other changes in emotional states of rats. Our data demonstrated that all phytocannabinoids reduced voluntary alcohol consumption in rats. However, compounds differed in their effectiveness in reducing alcohol consumption and the side-effect profile. Treatment with CBN and THCV reduced alcohol intake and demonstrated a good safety profile. In contrast, CBD only had a minor effect on alcohol consumption, reduced the locomotor activity and lowered the positive emotional states of rats. None of the compounds caused discomfort or distress. We conclude that phytocannabinoids acting as CB1 receptor partial agonists or neutral antagonists may have potential in treatment of AUD.

## INTRODUCTION

The endogenous cannabinoid system is the most abundant system in the brain, modulating nearly all physiological processes including mood, pain perception, sleep-wake cycle and appetite (for a review see Kano et al., 2009; Ligresti, et al., 2016; Aizpurua-Olaizola et al., 2017). Accordingly, it has been demonstrated that this system may be involved in many known pathological conditions, including several neurological and neurodegenerative disorders, as well as many psychiatric disorders like anxiety, depression and post-traumatic stress disorder (Ligresti et al., 2009; 2016; Basavarajappa, et al., 2017). It has also been established that the endocannabinoid system is involved in the development of substance use disorders (SUD, Maldonado et al., 2006). Furthermore, multiple preclinical studies demonstrated that reduction of CB1 receptor signalling was effective in reducing substance self-administration and relapse-related behaviours (Maldonado et al., 2006; Vengeliene et al., 2008), suggesting a good rationale for targeting the CB1 receptor to treat SUD. Hence, several CB1 antagonists/inverse agonists have been tested clinically; however, due to their adverse effects, they have not been found to be a suitable treatment option for patients suffering from SUD (Robinson et al., 2018; Nguyen et al., 2019).

Cannabis plants have long been used both medicinally and recreationally, mainly due to the psychoactive compound delta9-tetrahydrocannabinol (THC, a partial agonist of the CB1 receptor). However, health benefits of these plants may be attributable to over a hundred of other non-psychoactive compounds or their metabolites, collectively termed phytocannabinoids (Gülck and Møller, 2020). Phytocannabinoids act on the CB1 receptor either as neutral antagonists, partial agonists or allosteric modulators (Gülck and Møller, 2020), and may have a better safety profile than CB1 antagonists/inverse agonists that suppress constitutive receptor activity (e.g., Manning et al., 2021). The full therapeutic potential of phytocannabinoids is not yet known, and the majority of studies have focused on studying a major phytocannabinoid, cannabidiol (CBD, a negative allosteric modulator of the CB1 receptor). Besides the cannabinoidergic system, CBD affects several other brain systems: opioidergic, serotonergic, TRPV channels and others (Pertwee, 2008; McPartland et al., 2015). It has been demonstrated that CBD has the potential to treat a variety of medical conditions, such as anxiety, epilepsy and chronic pain (Mechoulam et al., 2002).

Targeting the endocannabinoid system by use of non-psychoactive phytocannabinoids have recently been suggested for clinical development of SUD pharmacotherapies (e.g., Spanagel, 2020; Karimi-Haghighi et al., 2022). For instance, preclinical studies demonstrated that CBD may be protective against binge ethanol-induced neurotoxicity (Liput et al., 2013; Hamelink et al., 2008), alleviate nicotine withdrawal symptoms (Smith et al., 2021), attenuate cognitive deficits and neuroinflammation induced by early alcohol exposure (García-Baos et al., 2021) and reduce alcohol seeking and self-administration (Turna et al., 2019). Unfortunately, results of recent clinical trials using CBD to treat alcohol use disorder (AUD) did not yield promising results (Kirkland et al., 2025; Mueller et al., 2025), suggesting that more research is needed to evaluate therapeutic potential of CBD and other phytocannabinoids in treating SUD.

As mentioned above, despite growing interest in use of phytocannabinoids as potentially new therapeutic options for SUD, the majority of studies have been focused on the widely known major cannabinoids CBD and THC. Hence, in the present study we explored, for the first time, the impact of cannabinol (CBN) and tetrahydrocannabivarin (THCV) on voluntary alcohol consumption in long-term drinking rats. We also tested CBD under the same experimental conditions as a positive control experiment, since it has already been demonstrated that this compound reduced alcohol consumption in rodents (Viudez-Martínez et al., 2018). CBN is a partial agonist of the CB1 receptor, it activates this receptor but with much lower potency than the psychoactive compound THC (Rhee et al., 1997). CBN also acts at several other receptors, e.g. CB2 and TRPV channels (Rhee et al., 1997; De Petrocellis et al., 2011). CBN is mildly sedative and may have potential in treating sleep disorders (Lavender et al., 2023; Bonn-Miller et al., 2024; Arnold et al., 2025). Preclinical studies suggest that restoration of the circadian activity and sleep quality in chronically drinking animals could contribute to reduced likelihood of alcohol consumption (Lawrence et al., 2006; Vengeliene et al, 2020). THCV is a neutral antagonist of the CB1 receptor, however, at higher doses it may behave as “indirect agonist” by inhibiting elimination of endocannabinoids. THCV also acts at several other receptors, such as CB2, TRPV channels and 5-HT1A (Pertwee 2008; McPartland et al. 2015). It has been suggested that THCV and other neutral antagonists may have potential in treating obesity and SUD (Le Foll et al., 2009; Abioye et al., 2020; Mendoza, 2025).

To achieve the study goals, we used a long-term voluntary alcohol drinking model in male Wistar rats. This model is used to study changes in the maintenance of drinking behaviour and has good predictive validity (i.e., voluntary drinking in rats has been reduced by clinically used drugs acamprosate and naltrexone, e.g., Cowen et al., 2005; Waeiss et al., 2019). We also explored the side-effect profile of CBN, THCV and CBD by measuring treatment-induced changes in water consumption, body weight and home-cage locomotor activity of long-term voluntary alcohol drinking rats. To measure if compounds caused a state of discomfort or distress and other changes in emotional states of rats, we recorded ultrasonic vocalisations (USVs) in alcohol naïve group-housed rats. Three main subtypes of 50 kHz vocalizations expressing positive emotional states were counted: contact calls (simple USVs), calls emitted in highly emotional situations expressing arousal state (frequency-modulated calls with trills or just trills) and calls emitted in rewarding situations (step calls, i.e., frequency-modulated calls without trills). In addition, two subtypes of 22 kHz vocalizations were counted that are either emitted during exposure to a dangerous situation (long calls) or express a state of discomfort or distress (short calls) (Wright et al., 2010; Brudzynski, 2015; Poceviciute et al. 2023). Alcohol naïve grouped-housed rats were used, as our pilot studies showed that long-term alcohol drinking single-housed rats emit few, or no USVs at all.

## MATERIALS AND METHODS

### Animals

Fifty-one eight-week-old outbred male Wistar rats were used in the alcohol consumption study and 32 eight-week-old outbred male Wistar rats were used for studies in alcohol naïve rats (from our own breeding colony at the Vilnius University, Lithuania). All animals were housed in groups throughout their adolescence into adulthood. For the alcohol study, adult animals were housed individually in standard rat cages and for the vocalisation recordings all animals remained grouped-housed throughout the entire course of the experiment. Animals were kept under a 12/12-hour artificial light/dark cycle (lights on at 7:00 a.m.) and constant room temperature (22±1°C). Standard laboratory rat food (4RF21-GLP, Mucedola srl, Milan, Italy) and tap water were provided *ad libitum* throughout the experimental period. All experimental procedures were approved by the State Food and Veterinary Service of the Republic of Lithuania and were carried out in accordance with the local Animal Welfare Act and the European Communities Council Directive of 22 September 2010 (2010/63/EU).

### Drugs

For repeated administration of phytocannabinoids in the alcohol consumption study, CBN and THCV (generously provided by Sanobiotec, Lithuania) were dissolved in tween 80 (Sigma-Aldrich, Germany) and then diluted with 0.9% saline to a final tween concentration of 4% and 5%, respectively, and injected intraperitoneally (IP) as a volume of 2 ml/kg. CBD (THC Pharm GmbH, Germany) was suspended in 0.9% saline containing 0.5% methylcellulose (Sigma-Aldrich, Germany) and injected as a volume of 3 ml/kg IP. For a single administration of phytocannabinoids in alcohol naïve rats, all compounds were dissolved in tween 80 and then diluted with 0.9% saline to a final tween concentration of 20% and injected IP as a volume of 2 ml/kg (please note that 20% tween was used in order to dissolve CBD and ensure its fast absorption, as the USV recordings lasted for 5 minutes). Control animals received an equal volume of respective vehicles.

### Effect of phytocannabinoid treatment on alcohol consumption and locomotor activity

#### Voluntary alcohol consumption

After two weeks of habituation to the experimental room, all rats were given *ad libitum* access to water and to 5% ethanol solution (v/v) for one week, afterwards alcohol concentration was increased to 8% (v/v) for another week, and finally, it was increased to 10% (v/v) for the remainder of the experiment. Alcohol drinking solutions were prepared from the absolute (99.8%) ethanol (Honeywell, Germany) and then diluted with tap water. The positions of bottles were changed weekly to avoid location preferences. Water and alcohol intake were measured either weekly or daily, and from these data, alcohol consumption (in g of pure ethanol/kg of body weight per day, g/kg/day) and water consumption (in ml/kg/day) were calculated.

#### Locomotor activity recordings

To measure the effect of treatment on locomotor activity, rats were placed in rectangular acrylic cages (floor: 40 × 40 cm, height: 50 cm) to ensure enough moving space and to avoid recording interruptions caused by a rat moving under the bottle/food area. Cages were equipped with ports for food and 2 bottles (placed on the outer surface of the cage), and rats were given *ad libitum* access to food, tap water and to 10% ethanol solution. Passive infrared (PIR) sensors (SEN0171, DFRobot, Shanghai, China) were used for measuring 24-hour locomotor activity of rats. A sensor was activated by infrared light radiating from moving objects in its field of view. Each PIR sensor was placed inside of a 3D printed casing, placed above the cage and had a 60° detection angle. The sensors were powered by and connected to the digital pins of an Arduino Uno R3 microcontroller, programmed to send a timestamp and pin ID whenever a change of state (ON or OFF) was detected (the microcontroller code is available at https://github.com/MartynasArba/motionsensors). Since the Arduino Uno R3 has no clock module, the output was set to display milliseconds from turning the device on. This output was sent to a computer via a serial port, for which the baud rate was set to 9600. Data was captured using the Realterm 3.0.1.44 terminal program, which allowed writing the output to a.txt file with an added system timestamp. To evaluate the locomotor activity of rats, the sum movement duration was calculated by subtracting the time of every ON event (when motion was detected) from the time of the subsequent OFF event.

#### Pharmacological intervention study

In order to study the effect of phytocannabinoid treatment on baseline alcohol and water drinking, rats were divided into three groups (n=12 per treatment condition) in such a way that the mean baseline total alcohol and water intake were matched, and ensuring random assignment to the groups for each subsequent treatment to avoid carry-over effects from the previous treatment. Only those rats that had an average baseline alcohol intake of approximately 2 g/kg/day or more were used for the pharmacological intervention study. Before each drug treatment, stable baseline drinking was monitored for at least three days. Locomotor activity was measured in a subset of animals (n=6 per treatment condition) starting at least three days before drug treatment, during treatment days and for several post-treatment days.

Pharmacological intervention was done approximately once per month starting at the 6^th^ month of continuous alcohol consumption. First, after the last day of baseline measurements, each animal was subjected to a total of three daily injections of vehicle, 5 mg/kg of CBN or 10 mg/kg of CBN, given at approximately 6 p.m. Thereafter, animals were left undisturbed until baseline drinking levels recovered. As it was found that CBN produced a delayed effect on drinking, this treatment was repeated using either vehicle or 5 mg/kg of CBN injected at the beginning of the light phase (approximately 8 a.m.) to examine if delayed effect of CBN is related to its accumulation in the body or acute pharmacological effect of CBN is necessary to induce a long-lasting change in animal behaviour. Similarly, with one-month of the recovery period in between treatment periods, vehicle, 5 mg/kg of THCV or 20 mg/kg of THCV, and later vehicle, 20 mg/kg of CBD or 40 mg/kg of CBD, were administered as three daily injections at approximately 6 p.m. THCV and CBD were administered just before the animal’s active phase. The doses were chosen based on published reports and our pilot studies. Body weights were measured just before pharmacological intervention experiments and the next day after the last drug administration. The person responsible for animal handling was blind to the treatment assignment throughout the experiment.

### Effect of phytocannabinoid treatment on ultrasonic vocalisations in alcohol naïve rats

During USV recordings, rats were approximately 7-month-old. Testing took place in Eurostandard type IVS cage placed in a sound-attenuated cubicle. For acoustic data acquisition, an electret ultrasound microphone with a frequency response range of 0 to 125 kHz (Avisoft Bioacoustics, Germany) was securely positioned through a small opening on the top of the cubicle. All rats were habituated to the testing conditions 4 times: 3 times (every second-third day) by placing all cage-mates together into the testing cage and once individually (one day prior to the test) for 5 min. For acoustic data acquisition, each rat was separated from the group and gently placed into the testing cage for 5 min. Acoustic data acquisition was done using the software Avisoft-SASLab Pro (Avisoft Bioacoustics, Germany) under baseline conditions and repeated in approximately 2 weeks following either vehicle, 5 mg/kg of CBN, 5 mg/kg of THCV or 20 mg/kg of CBD (n=8 per treatment condition) administration 30 min prior to the test (please note that these doses were chosen as they demonstrated better safety profile than higher doses in long-term alcohol drinking rats). Rats were divided into groups so that the mean baseline number of different USV types was matched. All calls were manually selected from spectrograms by a trained observer blind to the experimental design. USVs were classified into three major subtypes of 50 kHz USVs (i.e., vocalizations expressing positive emotional states): (1) simple calls, (2) trills and frequency-modulated calls with attached trills and (3) all types of step calls; as well as 2 subtypes of 22 kHz USVs (i.e., vocalizations expressing negative emotional states): (1) long calls and (2) short calls.

### Statistics and data analysis

Data derived from baseline home-cage drinking (alcohol and water intake) and locomotor activity (for data analysis the time spent moving during the active (dark) phase was used) was analysed using a two-way analysis of variance (ANOVA) with repeated measures (factors were: treatment group and time (days)). Data analysis regarding the effects of treatment on the change in the animals’ body weight (%) was performed using a one-way ANOVA (factor – treatment group). Data derived from acoustic data acquisition was analyzed using a two-way ANOVA with repeated measures (factors were: treatment group and session (baseline vs. treatment)). Whenever significant differences were found, post-hoc Student Newman Keuls tests were performed. The chosen level of significance was p < 0.05.

## RESULTS

### Effect of phytocannabinoid treatment on alcohol consumption and locomotor activity

#### CBN treatment effect

A two-way repeated measures ANOVA revealed that treatment with CBN had a significant effect on alcohol drinking in rats [factor treatment group × time interaction effect: F(16,323)=2.76, p<0.001]. Post hoc analysis demonstrated that treatment with CBN caused a significant reduction of alcohol consumption compared to vehicle treatment and reduced alcohol intake below baseline drinking levels (Fig. 1A). Lower alcohol intake in rats treated with CBN persisted for 3 post-treatment days. Water intake data analysis demonstrated that CBN treatment caused a long-lasting increase in water intake compared to both vehicle treatment and baseline water consumption [factor treatment group × day interaction effect: F(16,323)=3.07, p<0.001] (Fig. 1B), demonstrating that this treatment was selective towards lowering alcohol consumption. Post hoc analysis revealed that the strongest effect on water intake was seen during post-treatment days, suggesting that CBN might be accumulating in the body. However, 5 mg/kg of CBN administered on 3 consecutive days during the onset of the rat’s inactive phase had no effect on either alcohol or water consumption (data not shown), suggesting that the acute pharmacological effect of CBN is necessary to lower alcohol consumption in rats.

**Figure 1.**
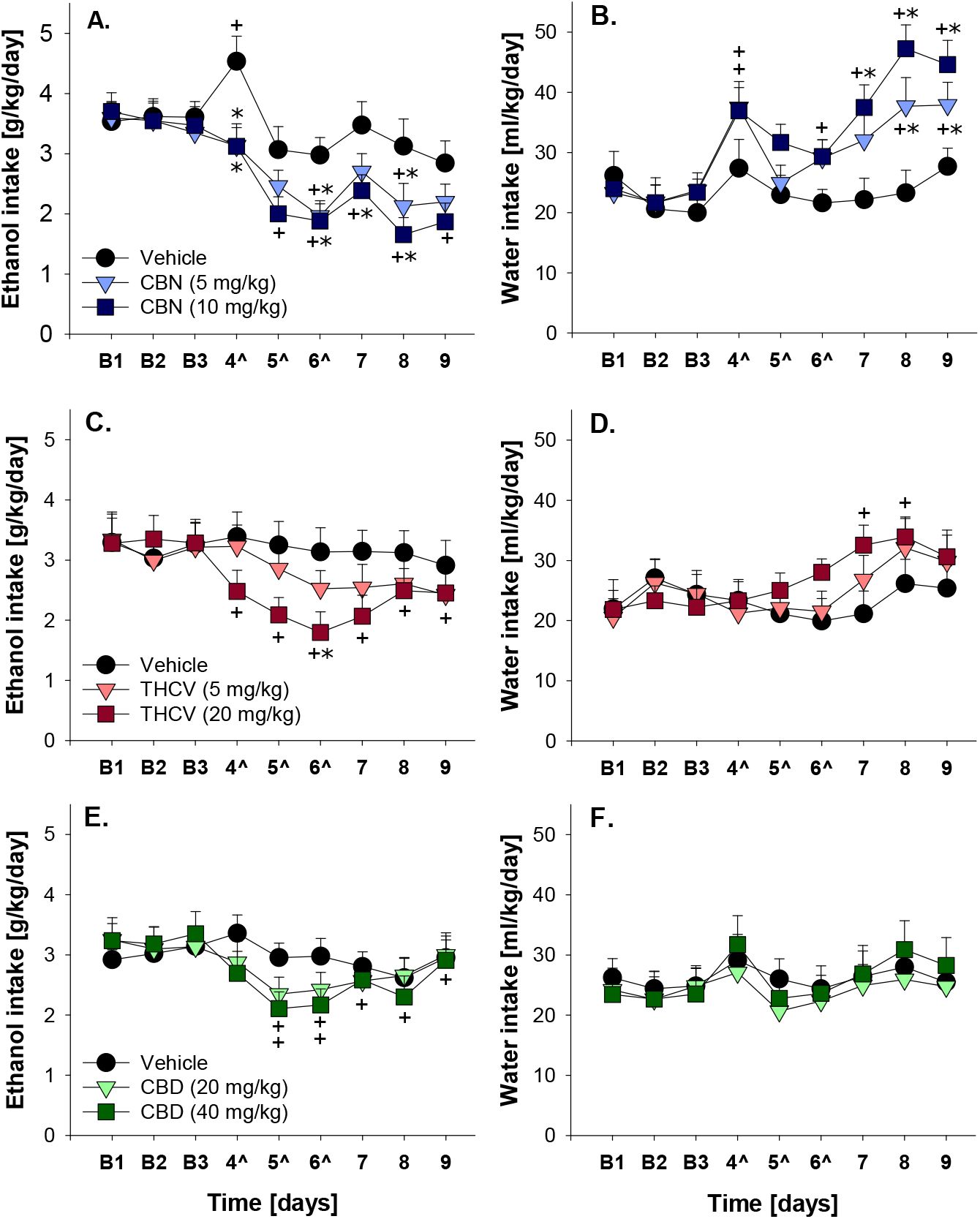
Total daily alcohol (calculated in g of pure ethanol per kg of body weight per day, g/kg/day, A, C, E) and water (ml/kg/day, B, D, F) intake in long-term drinking rats measured prior to (days B1-B3), during (days 4^-6^) and after (days 7-9) three once every 24 h administrations (just before the animal’s active phase) of (A, B) vehicle, 5 mg/kg of CBN or 10 mg/kg of CBN; (C, D) vehicle, 5 mg/kg of THCV or 20 mg/kg of THCV; and (E, F) vehicle, 20 mg/kg of CBD or 40 mg/kg of CBD (n=12 per treatment condition). Data are presented as means ± S.E.M. * indicates significant differences from the vehicle group and + indicates significant difference from all 3 baseline days, p<0.05.

Analysis of the locomotor activity data by use of a two-way repeated measures ANOVA revealed that treatment with CBN slightly but not significantly reduced the time spend moving during the active phase when compared to vehicle treatment [factor treatment group: p=0.07 and factor treatment group × time interaction effect: p=0.26] (Fig. 2A). Lower dose of CBN did not have a significant impact on the body weight, however, treatment of rats with the higher dose of CBN caused small but significant loss (−1.3±0.4%) of the body weight [factor treatment group: F(2,23)=5.15, p<0.05]. These data suggest that CBN had a mild sedative effect on rats and 10 mg/kg dose affected food intake and/or metabolism.

**Figure 2.**
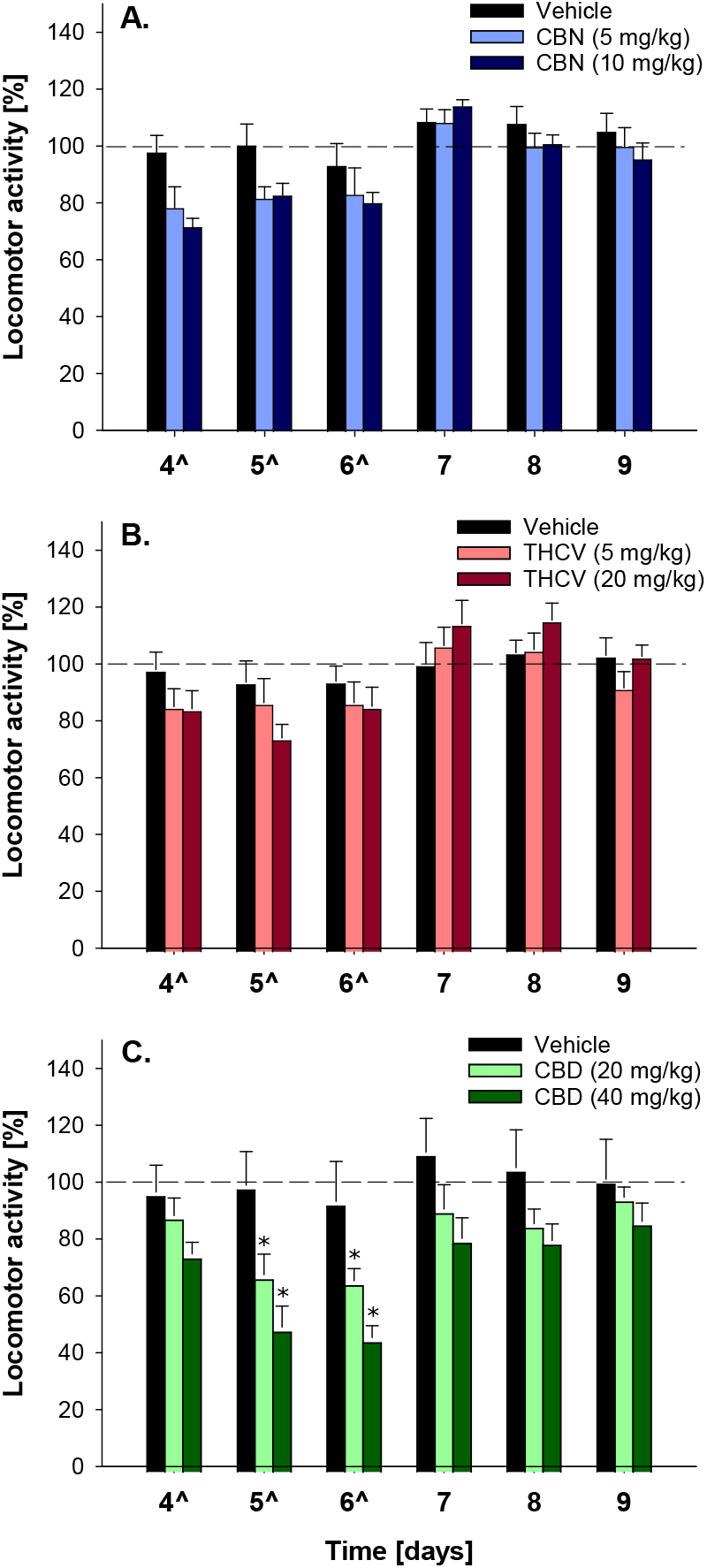
Locomotor activity in (A) vehicle, 5 mg/kg of CBN or 10 mg/kg of CBN, (B) vehicle, 5 mg/kg of THCV or 20 mg/kg of THCV, and (C) vehicle, 20 mg/kg of CBD or 40 mg/kg of CBD treated long-term drinking rat groups (n=6 per treatment condition). Locomotor activity is shown as shown as the time spent moving during 12-hour postinjection intervals of the animals’ active phase. The percentage of each rat’s locomotor activity during (days 4^-6^) and after (days 7-9) treatment was calculated with respect to basal activity prior to treatment (dashed line). Data are presented as means ± S.E.M. * indicates significant differences from the vehicle group, p<0.05.

#### THCV treatment effect

A two-way repeated measures ANOVA demonstrated that THCV treatment caused a significant reduction in alcohol intake compared to vehicle treatment and reduced alcohol intake below baseline drinking levels [factor treatment group × day interaction effect: F(16,323)=3.93, p<0.001]. However, a subsequent post hoc analysis showed that alcohol consumption was significantly different between vehicle and THCV treated rats only following the third administration of the higher THCV dose (Fig. 1C). Analysis the water consumption data revealed that THCV treatment also had a significant effect on water intake [factor treatment group × day interaction effect: F(16,323)=1.78, p<0.05]. Post hoc analysis revealed that 20 mg/kg of THCV increased water consumption above baseline levels during two post-treatment days (Fig. 1D).

Both locomotor activity and body weights were affected by repeated THCV administration. Sum time spent moving during the active phase was slightly lower during treatment days but recovered immediately as drug administration was terminated [factor treatment group: p=0.71 and factor treatment group × time interaction effect: F(10,107)=2.19, p<0.05] (Fig. 2B). Similarly to CBN, the higher dose of THCV caused small but significant loss (−1.2±0.3%) of the body weight [factor treatment group: F(2,23)=11.24, p<0.001].

#### CBD treatment effect

Drinking data analysis revealed a significant effect of CBD treatment on alcohol consumption [factor treatment group × day interaction effect: F(16,323)=2.24, p<0.01]. Post hoc analysis showed that this treatment reduced alcohol consumption below baseline drinking levels; however, differences in alcohol intake between treatment groups were not significant (Fig. 1E). Furthermore, water consumption was unaffected by this treatment [factor treatment group × day interaction effect: p=0.91] (Fig. 1F).

CBD treatment had significant effect on the locomotor activity of rats [factor treatment group: F(2,107)=4.39, p<0.05 and factor treatment group × time interaction effect: F(10,107)=2.64, p<0.01] (Fig. 2C). The higher dose of this treatment also caused a slight reduction (−0.5±0.5%) of the body weight of animals [factor treatment group: F(2,23)=4.33, p<0.05].

### Effect of phytocannabinoid treatment on ultrasonic vocalisations in alcohol naïve rats

A two-way repeated measures ANOVA revealed that administration of test compounds lowered the number of 50 kHz USVs compared to the baseline recording session prior to treatment [factor session: F(1,63)=9.64, p<0.01], and specifically, reduced the number of simple calls and trills [factor session: F(1,63)=6.33, p<0.01 and F(1,63)=12.05, p<0.01 for simple calls and trills, respectively]. Post hoc analysis revealed that this reduction was statistically significant only for the 20 mg/kg of CBD treatment group (Fig. 3). Neither compound reduced the number of 50 kHz USVs [factor treatment group × session interaction effect: p=0.51; p=0.51; p=0.75 and p=0.58 for sum 50 kHz calls, simple calls, trills and step calls, respectively] below that recorded in the vehicle treated group.

**Figure 3.**
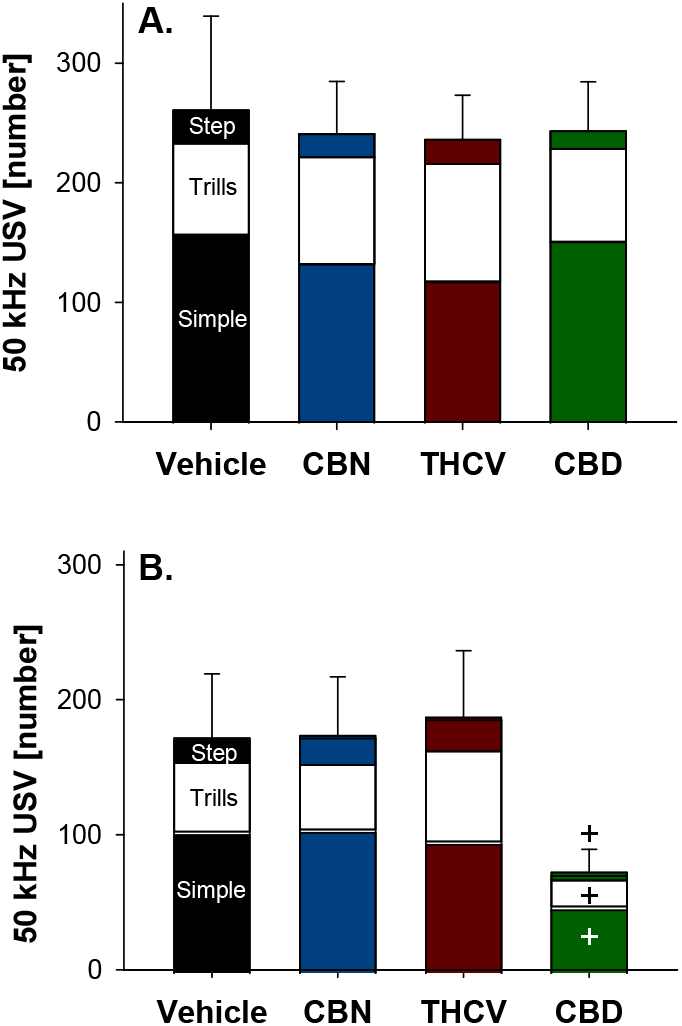
The number of 50 kHz ultrasonic vocalisations (USVs) in alcohol naïve male rats emitted during 5-min acoustic recordings (A) before and (B) after a single administration of vehicle, 5 mg/kg of CBN, 5 mg/kg of THCV or 20 mg/kg of CBD (n=8 per treatment condition), 30 min prior to the recording session. Three main subtypes of 50 kHz vocalizations expressing positive emotional states were quantified as simple calls, trills or calls with attached trills and all types of step calls without trills. There were no 22 kHz USVs during either baseline or treatment session. Data are presented as the mean number of simple calls, trills and step calls and as the mean number of 50 kHz USVs ± S.E.M. + indicates significant differences from the baseline condition, p < 0.05.

Rats did not emit any 22 kHz calls during either baseline or post-treatment recording sessions, demonstrating that a single administration of either 5 mg/kg of CBN, 5 mg/kg of THCV or 20 mg/kg of CBD did not cause discomfort or distress to the animals.

## DISCUSSION

Our study demonstrated that repeated administration of all three phytocannabinoids reduced voluntary alcohol intake in long-term drinking male Wistar rats. However, the compounds differed in their effectiveness and the side-effect profile. Administration of CBN caused a significant reduction in alcohol intake, and it was accompanied by a marked increase in water intake, demonstrating the selectiveness of this treatment towards lowering alcohol consumption. This effect was observed during three consecutive days following treatment completion. The long-lasting effect of CBN was not caused by its accumulation, since CBN administered during the onset of the rat’s inactive phase had no effect on either alcohol or water intake, suggesting that the acute pharmacological effect of CBN is necessary to induce a long-lasting change in animal behaviour. Treatment with THCV had similar but somewhat weaker effect on alcohol and water consumption. Finally, CBD had only a modest effect on alcohol consumption and did not affect water intake. CBN and THCV did not have a significant impact on the time spent moving during their active phase in chronically drinking animals. However, a marked reduction in the locomotor activity was seen during treatment with either dose of CBD. Higher doses of all compounds led to a small but significant decrease in the body weight, demonstrating that those (and higher) doses of these compounds may cause unselective effects in long-term voluntary alcohol drinking male rats. Lower doses of all three compounds had no effect on the body weight in chronically drinking rats and did not induce a state of discomfort or distress in alcohol naïve rats. However, the lower dose of CBD reduced expression of positive emotional state of rats as measured by the change in 50 kHz vocalizations. These data suggest that lower doses of CBN and THCV demonstrate a good safety profile and may be effective in lowering alcohol consumption, whereas any effective dose of CBD may cause unwanted effects.

It is known that endocannabinoid signalling contributes to the primary rewarding effects of alcohol due to its modulatory function on the mesocorticolimbic dopaminergic system; and CB1 receptor blockade attenuates voluntary alcohol intake and several other alcohol-related behaviours (Maldonado et al., 2006; Basavarajappa, 2007). Therefore, it is surprising that in our study, CBN, a low potency partial agonist, was more effective in lowering alcohol consumption than THCV or CBD. It has been demonstrated that chronic alcohol exposure may lead to increased extracellular levels of endocannabinoids and down-regulation of CB1 receptor expression and function (Basavarajappa, 2007). This suggests that under the present experimental conditions (rats had continuous access to alcohol for at least 6 months), endocannabinoid levels may have been elevated and CBN acted as an indirect CB1 receptor antagonist via interference with the endogenous agonist binding. Furthermore, it has been shown that endocannabinoid signalling may be reversed by alcohol abstinence (Pava and Woodward, 2012 Kleczkowska et al., 2016); hence, response to CB1 receptor partial agonists following short alcohol exposure or using abstinent rats may be different than that in chronically drinking rats. Indeed, it has been demonstrated that administration of partial agonist of the CB1 receptor THC promoted relapse-like behaviour in alcohol abstinent rats (McGregor et al., 2005), however, the study of McMillan and Snodgrass (1991) showed that acute administration of THC reduced maintenance of alcohol consumption in rats that had been trained to consume alcohol in an operant setting for two months.

It has been demonstrated that CB1 antagonists/inverse agonists were effective in both body weight reduction (Murphy and Le Foll, 2020) and self-administration of many abused substances (Maldonado et al., 2006). A neutral antagonist of the CB1 receptor THCV has also been effective in appetite suppression and increased energy expenditure (Mendoza, 2025), and thus, it could be expected that this compound would produce similar effect on alcohol consumption as CB1 antagonists/inverse agonists. Our study supports this notion, demonstrating that THCV dose-dependently reduced alcohol intake that was accompanied by a compensatory increase in water consumption.

The effectiveness of CBD, a negative allosteric modulator of the CB1 receptor, on alcohol-related behaviours has already been demonstrated in several preclinical studies. Both acute and repeated daily administration of CBD has been shown to reduce voluntary alcohol consumption, self-administration and had a long-lasting effect on relapse-related behaviours (Gonzalez-Cuevas et al., 2018; Viudez-Martínez et al., 2018; Maccioni et al., 2022). In our study, CBD had only a small and short-lasting effect on voluntary alcohol consumption. Increasing the dose of CBD may have had a stronger impact on alcohol consumption,;however, as mentioned above, the highest dose of CBD caused a significant loss of the body weight, indicating the occurrence of unwanted effects. In addition, both doses of CBD caused a marked reduction in home-cage activity. Compared to earlier studies, our rats were not only older (during CBD treatment they were approximately 1 year old) but also had been drinking alcohol for a long period of time. This could explain higher sensitivity of these rats to CBD with respect to unwanted effects. Taking into consideration recent clinical trials showing that CBD had no effect on alcohol drinking (Kirkland et al., 2025; Mueller et al., 2025) it is likely that CBD alone may not be sufficient to treat AUD without inducing unwanted effects (see also Redonnet et al, 2025).

In summary, the present study demonstrated that CBN was the most effective phytocannabinoid in reducing the maintenance of voluntary alcohol consumption in male rats when compared to either THCV or CBD. The effect of all three phytocannabinoids on alcohol consumption may be related to their action on the CB1 receptor. However, all three compounds have multiple other molecular targets that may have contributed to their effect on rat behaviour. Because of their different mechanisms of action, phytocannabinoids, such as those used in the present study, could be beneficial in treating different aspects of AUD (i.e., maintenance, withdrawal, or relapse prevention) and thus, contribute to the development of personalized treatment strategies. For instance, CBN may not be effective in relapse prevention; however, it might be effective as a substitution therapy in chronically drinking individuals. Finally, the anti-inflammatory and neuroprotective action of phytocannabinoids (Hamelink et al., 2008; Liput et al., 2013; Gojani et al., 2023) provides an additional beneficial effect in AUD treatment owing to their ability to reduce neurotoxic consequences of chronic alcohol consumption (Charlton et al., 2019).

## ACKNOWLEDGEMENTS

This work was supported by the Lithuanian Business Support Agency (S-01.2.1-LVPA-K-856-01-0222).

